# Inhibition of the SUV4-20 H1 histone methyltransferase increases frataxin expression in Friedreich’s ataxia patient cells

**DOI:** 10.1101/2020.03.26.010439

**Authors:** G Vilema-Enríquez, R Quinlan, P Kilfeather, R Mazzone, S Saqlain, I del Molino del Barrio, A Donato, G Corda, F Li, M Vedadi, AH Németh, PE Brennan, R Wade-Martins

## Abstract

The molecular mechanisms of reduced frataxin (*FXN*) expression in Friedreich’s ataxia (FRDA) are linked to epigenetic modification of the *FXN* locus caused by the disease-associated GAA expansion. Here, we identify that SUV4-20 histone methyltransferases, specifically SUV4-20 H1, play an important role in the regulation of *FXN* expression and represent a novel therapeutic target. Using a human *FXN*-GAA-Luciferase repeat expansion genomic DNA reporter model of FRDA, we screened the Structural Genomics Consortium epigenetic probe collection. We found that pharmacological inhibition of the SUV4-20 methyltransferases by the tool compound A-196 increased the expression of *FXN* by approximately 1.5-fold in the reporter cell line and in several FRDA cell lines and patient-derived primary peripheral blood mononuclear cells. SUV4-20 inhibition was accompanied by a reduction in H4K20me2 and H4K20me3 and an increase in H4K20me1, but only modest (1.4–7.8%) perturbation in genome-wide expression was observed. Finally, based on the structural activity relationship and crystal structure of A-196, novel small molecule A-196 analogues were synthesized and shown to give a 20-fold increase in potency for increasing *FXN* expression. Overall, our results suggest that histone methylation is important in the regulation of *FXN* expression, and highlight SUV4-20 H1 as a potential novel therapeutic target for FRDA.

## INTRODUCTION

Friedreich’s ataxia (FRDA) is the most common autosomal recessive ataxia with an estimated prevalence of 1 in 50,000 in the Caucasian population (1, 2). The disorder is caused by an unstable GAA trinucleotide repeat expansion in the first intron of the frataxin (*FXN*) gene locus (1) on chromosome 9 (3). *FXN* encodes frataxin, a protein which plays a role in mitochondrial iron-sulphur cluster biogenesis and is highly conserved across most organisms (4). The aberrant GAA expansion leads to partial transcriptional silencing of *FXN*, which results in the expression of structurally and functionally normal frataxin, but at dramatically lower levels compared to the wild-type locus (5). In the normal *FXN* gene there can be up to 40 GAA repeats, whereas disease-associated alleles contain more than 40 GAA repeats, most commonly around 600-900. Larger GAA expansions, particularly that of the smaller allele, correlate with earlier age at onset and severity of the disease (6).

The precise mechanism by which the GAA expansion causes a partial silencing of *FXN* is still unclear. However, a wealth of studies have documented that expanded GAA.TTC repeats adopt unusual DNA structures which are responsible for the reduced levels of frataxin. These unusual structures might produce either sticky DNA (formed by the association of two purine.purine.pyrimidine triplexes (7–11)), persistent DNA-RNA hybrids (12), or induce the formation of repressive heterochromatin (13, 14). Furthermore, gene silencing has been recently linked to its association with the nuclear transcriptional repressive environment, the nuclear lamina (15).

In recent years, epigenetic changes such as DNA methylation or histone modifications have been implicated in a variety of diseases including FRDA (16). The GAA triplet repeat expansion was shown to silence the *FXN* locus similar to position effect variegation (13), and be enriched in histone modifications associated with inactive heterochromatin (H3K9me2/3, H3K27me3, H4K20me3), and conversely to have reduced acetylated histones H3 and H4, which are marks of active chromatin (16, 17). Amongst the histone marks associated with silent chromatin, H4K20me3 is highly enriched at telomeres and pericentric heterochromatin, as well as imprinted regions and repetitive elements, suggesting that this histone modification is involved in transcriptional silencing (18). In FRDA, the *FXN* gene carrying a GAA expansion has been shown to have increased H4K20me3 in the flanking regions of the GAA repeats (19), suggesting that the transcriptionally repressive H4K20me3 may be involved in the silencing of *FXN*. The family of histone methyltransferases SUV4-20, comprising two lysine methyltransferase (KMT) enzymes SUV4-20 H1 and SUV4-20 H2, are responsible for the generation of H4K20me2 and H4K20me3 (20).

Here, we report that the inhibition of the SUV4-20 histone methyltransferases, specifically SUV4-20 H1, increases FXN protein expression in a human *FXN*-GAA-Luciferase repeat expansion genomic DNA locus reporter model and in primary FRDA patient-derived cells. This novel finding highlights the importance of the methylation of H4K20 in the silencing of *FXN* and identifies the SUV4-20 H1 methyltransferase as a novel target for therapeutic intervention.

## RESULTS

### Screening the Structural Genomics Consortium epigenetic probe collection identifies histone methyltransferases as important for the regulation of *FXN* expression

High levels of specific heterochromatin marks have been previously reported at the first intron of the pathologically silenced *FXN* gene (for a review of the epigenetic changes associated with *FXN* see (17)). The epigenetic probe collection from the Structural Genomics Consortium (SGC) comprises a well-characterised set of drug-like small molecules which inhibit specific chromatin regulatory proteins and domains including bromodomains, demethylases and methyltransferases (21, 22). To identify novel epigenetic targets involved in the regulation of *FXN* expression, we screened the SGC epigenetic chemical probe set using our *FXN*-GAA-Luc reporter cell line (15, 23) in a 96-well format in duplicate (see Supplementary Table 1) and assayed frataxin-luciferase (FXN-Luc) protein expression by luciferase assay (Figure 1A). The screen identified five candidate compounds able to increase the expression of FXN-Luc protein above the levels of the dimethyl sulphoxide (DMSO) vehicle control (Figure 1B) in the absence of any toxicity assayed by the adenylate kinase assay (Supplementary Figure 1). The positive hit compounds are inhibitors of several histone methyltransferases, being SUV4-20 H1/H2 (compound 3; A-196), G9a/GLP (compound 4; A-366), EZH1/EZH2 (compound 11; GSK343), type I protein arginine methyltransferases (PRMTs) (compound 23; MS023) and DOT1L (compound 33; SGC0946).

**Figure 1.**
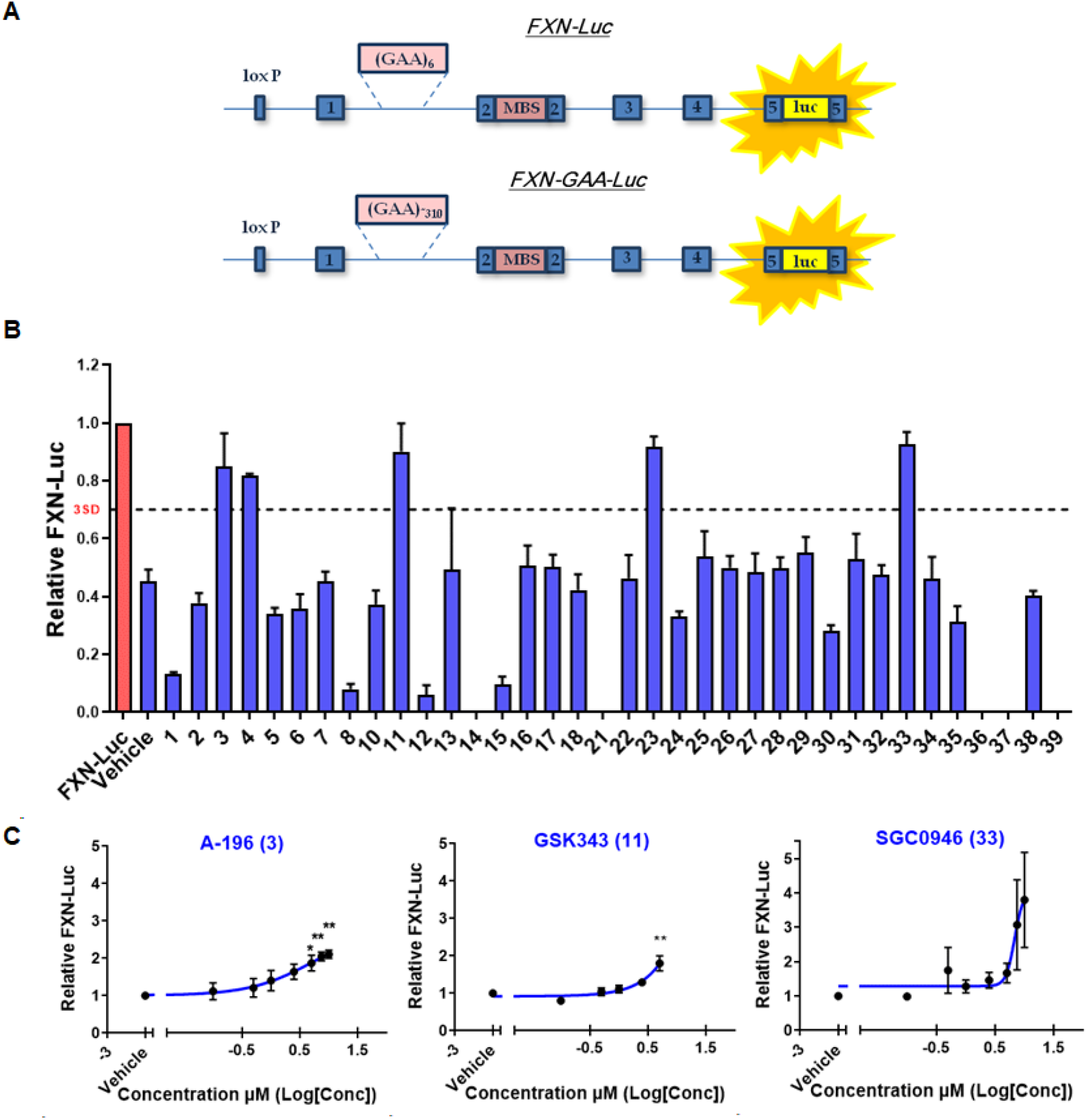
Screening the SGC epigenetic probe set identifies histone methyltransferases as regulators of *FXN* repression. **(A)** Schematic representation of the *FXN*-Luc and *FXN*-GAA-Luc reporter cell lines. **(B)** Luciferase assay of the *FXN*-GAA-Luc cell line treated with the SGC epigenetic probes collection. The *FXN*-Luc cell line was used as a reference for FXN levels. **(C)** Concentration-response curves assessed by luciferase assay of the *FXN*-GAA-Luc cell line treated for six days with A-196 EC_50_ 5.2 μM (left panel), GSK343 EC_50_ 596 μM (middle panel), and SGC0946 EC_50_ 6.8 μM (right panel). Data are relative to the vehicle and presented as mean ± SEM, n=3 performed in duplicate, one-way ANOVA followed by Bonferroni test *p<0.05, **p<0.01

We next performed a concentration-response assay for the five hits identified in the primary screen to study the expression of *FXN*-GAA-Luc protein with increasing concentration of each chemical probe. We incubated the *FXN*-GAA-Luc cell line in a 96-well format in triplicate with increasing concentrations of the probes from 0.1 μM to 10 μM for six days. Three out of the five probes were confirmed as hits and concentration-response curves were obtained (Figure 1C). Four of the chemical probes exhibited no toxicity, whereas GSK343 was toxic to cells at concentrations above 5 μM (see Supplementary Figure 1) and so *FXN*-GAA-Luc protein expression was only assessed from 0.1 μM to 5 μM of GSK343. EC_50_ values were estimated as 5.2 μM (compound 3), 596 μM (compound 11) and 6.8 μM (compound 33).

Overall, the screening of the SGC epigenetic probe collection demonstrated that inhibition of several methyltransferases acting on histones H3 and H4 may play an important role in the regulation of *FXN* expression. Furthermore, this work suggests histone methyltransferases as a novel set of potential therapeutic targets not previously studied for FRDA.

### Genetic modification of the SUV4-20 family of methyltransferases reveals histone H4 lysine 20 (H4K20) methylation as a key epigenetic mark for *FXN* gene silencing

The screen of the SGC epigenetic probe collection highlighted the role of histone lysine methylation in *FXN* silencing. Previous reports have described the accumulation of methylation marks on lysine residues on histones H3 and H4 at the *FXN* gene (13, 19, 24–31). To validate the most promising targets, we performed siRNA-mediated knockdown of *SUV4-20H1/H2*, *EZH1/EZH2* and *DOT1L* in the *FXN*-GAA-Luc cell line. *FXN*-GAA-Luc HEK293 cells were treated with 25 nM of siRNA for six days. siRNA-mediated knockdown of *SUV4-20 H1*, but not *SUV4-20 H2*, significantly increased FXN-Luc protein expression (Figure 2A-D) providing genetic validation of SUV4-20 H1 as the target of interest.

**Figure 2.**
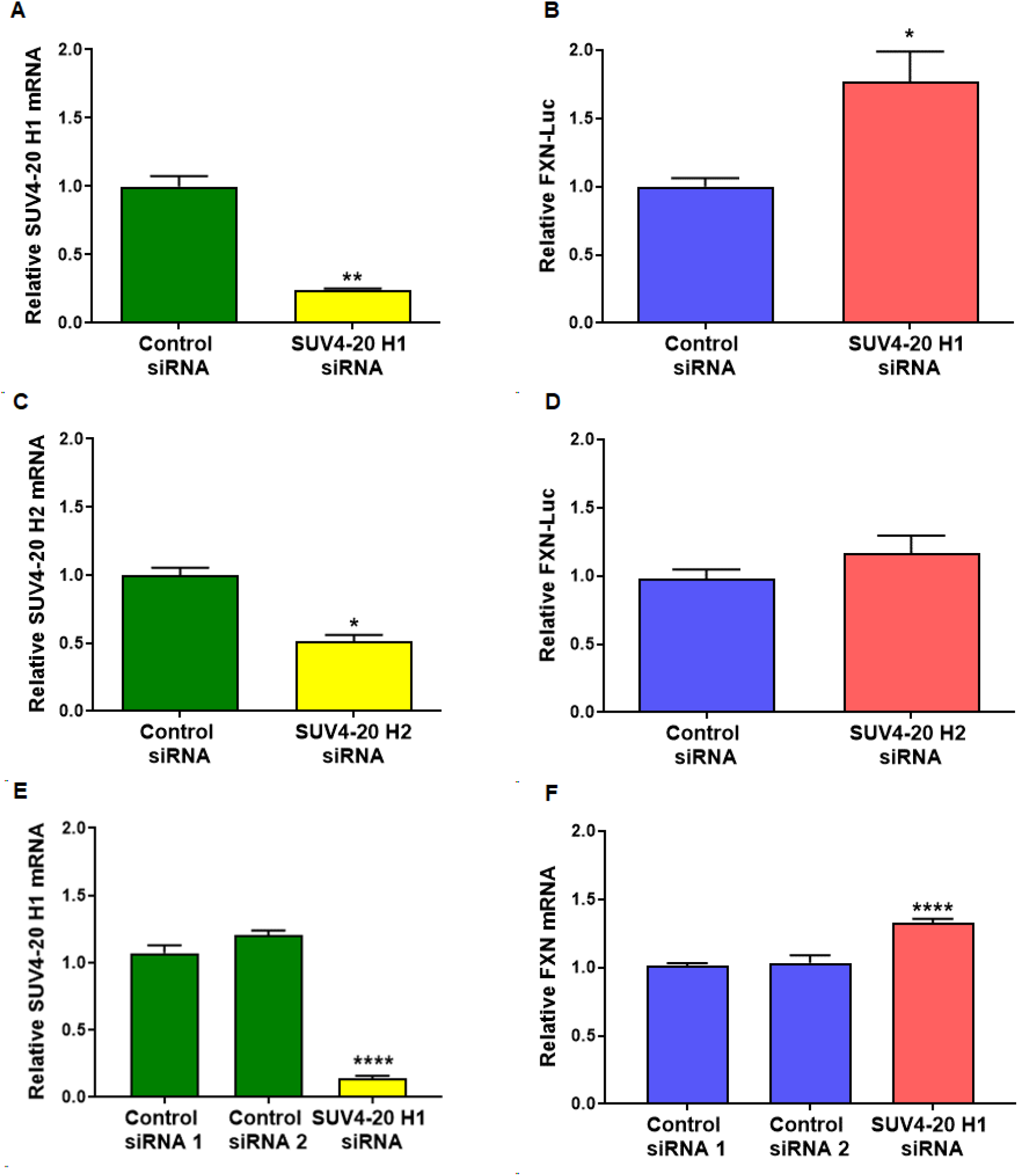
siRNA knockdown of the SUV4-20 family of methyltransferases identifies SUV4-20 H1 as a critical protein for the repression of *FXN*. **(A)** Relative *SUV4-20 H1* mRNA expression after *SUV4-20 H1* siRNA-mediated knockdown assessed by qRT-PCR. **(B)** Relative *FXN* mRNA expression after *SUV4-20 H1* siRNA-mediated knockdown assessed by qRT-PCR. **(C)** Relative *SUV4-20 H2* mRNA expression after *SUV4-20 H2* siRNA-mediated knockdown assessed by qRT-PCR **(D)** Relative *FXN* mRNA expression after *SUV4-20 H2* siRNA-mediated knockdown assessed by qRT-PCR. **(E)** Relative *SUV4-20 H1* mRNA expression after *SUV4-20 H1* knockdown assessed in the fibroblast line GM04078. **(F)** Relative *FXN* mRNA expression after *SUV4-20H1* down-regulation in fibroblasts GM04078. Experiments performed in the line GM04078 were carried out using two different control siRNA. Data are relative to control siRNA, and control siRNA 1, correspond to 6 days treatments and are presented as mean ± SEM, n=3 performed in triplicate, one-way ANOVA followed by Bonferroni test *p<0.05, **p<0.01, ***p<0.001.

However, siRNA-mediated knockdown of *DOT1L*, *EZH1* or *EZH2* did not result in a significant increase of FXN-Luc protein expression (Supplementary Figure 2, 3).

We then performed siRNA-mediated knockdown of *SUV4-20 H1* in the FRDA patient-derived primary fibroblast line GM04078 to confirm that the up-regulation of *FXN* following knockdown of SUV4-20 H1 is not limited to the *FXN*-GAA-Luc reporter cell line. The siRNA-mediated knockdown of *SUV4-20 H1* in primary fibroblasts increased *FXN* mRNA expression by approximately 1.25-fold (Figure 2E, F). Taken together, these results demonstrate that direct downregulation of SUV4-20 H1 significantly increases *FXN* expression, validating this histone lysine methyltransferase as a therapeutic target for FRDA.

### A-196 increases frataxin protein expression in FRDA patient-derived primary cells

To extend the finding that A-196 is able to increase *FXN* expression to FRDA patient cells we treated several patient-derived cells with 5 μM and 10 μM A-196 for six days. We first treated the primary fibroblast line GM04078 and assessed mature frataxin protein expression by western blot (Figure 3A, B). Treatment with A-196 significantly increased mature FXN expression at both concentrations in this primary fibroblast line.

**Figure 3.**
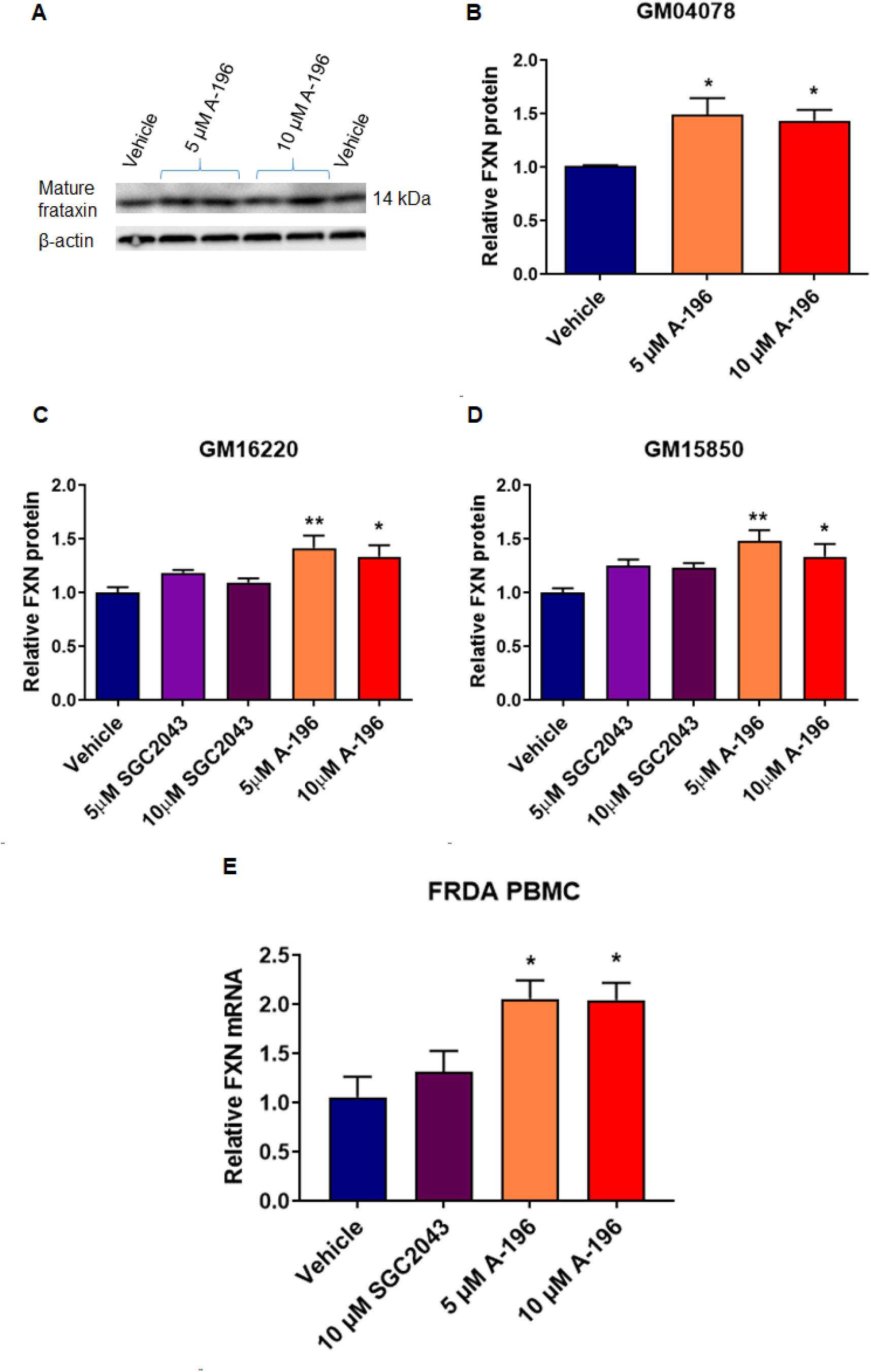
A-196 increases frataxin protein expression in patient-derived cells. **(A)** Representative western blot of mature frataxin protein expression in the primary fibroblast line GM04078 after A-196 treatment, n=3 in duplicates **(B)** Quantification of the experiment shown in panel A, **(C, D)** Relative frataxin protein expression in the lymphoblastoid cell lines GM16220 and GM15850 assessed by AlphaLISA, n=4 in triplicate **(E)** Frataxin protein expression after A-196 treatment in PBMCs extracted from four FRDA patients. Data are relative to the vehicle, a treatment of 6 days and are presented as mean ± SEM, one-way ANOVA followed by Bonferroni test *p<0.05, **p<0.01.

We next treated the FRDA patient-derived lymphoblastoid cell lines GM16220 and GM15850 with 5 μM and 10 μM A-196 for six days and measured total frataxin levels by AlphaLISA. As previously shown in fibroblasts, A-196 was able to increase significantly FXN protein expression in both lymphoblastoid cell lines (Figure 3C, D). We also showed that the structural inactive analogue of A-196, SGC2043, did not increase frataxin protein expression, further confirming the selective and specific mechanism of action of A-196.

Finally, we tested A-196 in primary peripheral blood mononuclear cells (PBMCs) isolated from FRDA patients. PBMCs treated with 5 μM and 10 μM A-196 for six days increased *FXN* mRNA expression by approximately 2-fold assessed by qRT-PCR (Figure 3E). Overall, these results confirm that the specific inhibition of the SUV4-20 methyltransferases increases frataxin levels in multiple FRDA patient-derived cells.

### Pharmacological inhibition of SUV4-20 decreased H4K20me2/3 and increased H4K20me1 in FRDA patient-derived cells

The mechanism of action of the highly selective histone methyltransferase inhibitor A-196 has been previously described (32) showing that A-196 decreases global levels of the repressive H4K20me2/3 mark while increasing H4K20me1. To better understand how the inhibition of SUV4-20 mediates the up-regulation of *FXN*, we first analysed the global methylation status of H4K20 in the *FXN*-GAA-Luc cell line after A-196 treatment. In addition to the structural inactive analogue SGC2043, we also tested another inactive analogue, A-197. As expected, treatment with 5 μM or 10 μM of A-196, but not with SGC2043 or A-197, decreases H4K20me2/3 with a concomitant increase in H4K20me1 (Figure 4A-D).

**Figure 4.**
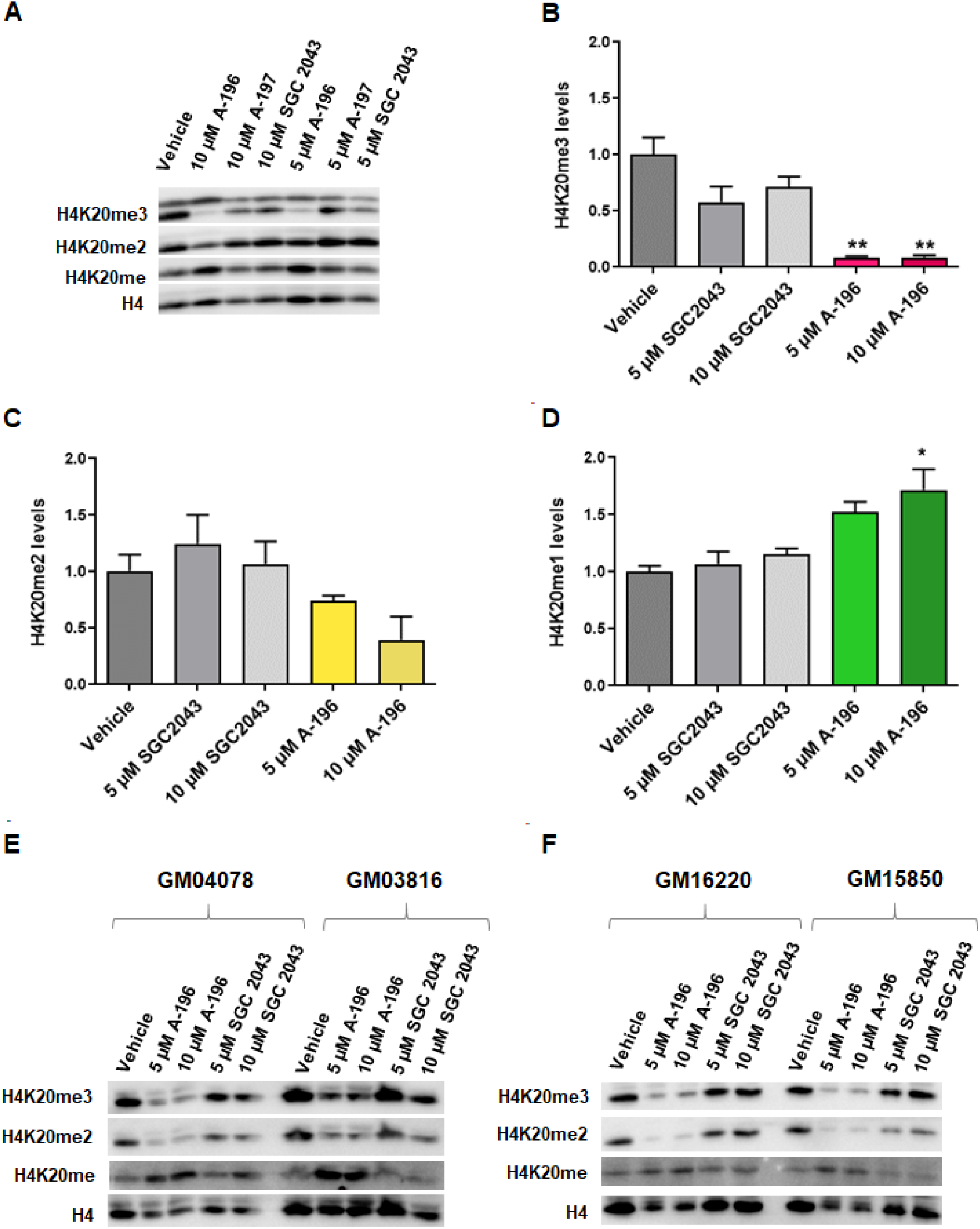
Pharmacological inhibition of the SUV4-20 methyltransferases with A-196 decreases H4K20me2/3 in the *FXN*-GAA-Luc cell line and in FRDA patient-derived cells. **(A)** Representative western blot of the *FXN*-GAA-Luc cell line after A-196, A-197 and SGC2043 treatment. **(B-D)** Quantification of the global level of H4K20 methylation after inhibition of SUV4-20. **(E and F)** Representative western blots of patient-derived cells after treatment with the above-mentioned probes. Data are relative to the vehicle, a treatment of 6 days and are presented as mean ± SEM, n=3, one-way ANOVA followed by Bonferroni test *p<0.05, **p<0.01.

We also tested the effect of A-196 on the H4K20me2/3 and H4K20me1 marks in FRDA patient-derived primary fibroblasts and lymphoblastoid cell lines compared to the inactive control SGC2043. A six-day treatment with 5 μM or 10 μM A-196, but not SGC2043, reduced the global levels of H4K20me2/3 and increased the levels of H4K20me1 (Figure 4E, F).

These results provide evidence that the inhibition of SUV4-20 by A-196 reduces global levels of the repressive H4K20me2/3 mark, increases the levels of H4K20me1, and increases *FXN* expression. When levels of H4K20me2/3 are unchanged, as in the case of the treatment with inactive probes, neither H4K20me2/me3, H4K20me1 or frataxin levels change (Figures 3 and 4). These results highlight the importance of H4K20 methylation in the regulation of expression of *FXN* and identify H4K20 methylation as an important histone post-translational modification for FRDA.

### Transcriptional perturbation after treatment with A-196 was concentration-dependent and limited to between 1% and 8% of protein-coding genes

Although the inhibition of SUV4-20 results in an increase in *FXN* expression which has potential therapeutic implications for FRDA, this inhibition may alter the epigenome at other loci, resulting in aberrant gene expression elsewhere. Therefore, we treated primary fibroblasts with 1 μM, 5 μM and 10 μM A-196 for six days and analysed genome-wide perturbations in gene expression due to the inhibition of SUV4-20 by RNA-Seq. We first assessed *FXN* mRNA expression, confirming an increase in expression (Figure 5A). We then performed principal component analysis (PCA) on the RNA-Seq data and determined that samples treated with A-196 show a concentration-dependent separation along PC1 from untreated, vehicle and inactive analogue-treated samples (Figure 5B). Next, we filtered genes with low read counts and performed differential gene expression analysis to quantify the extent of transcriptional perturbation. The lowest concentration of A-196 is responsible for 193 differentially-expressed genes (DEGs, out of ~14,000 genes measured), 5 μM A-196 for 626 DEGs and 10 μM A-196 for 1098 DEGs (Figure 5C).

**Figure 5.**
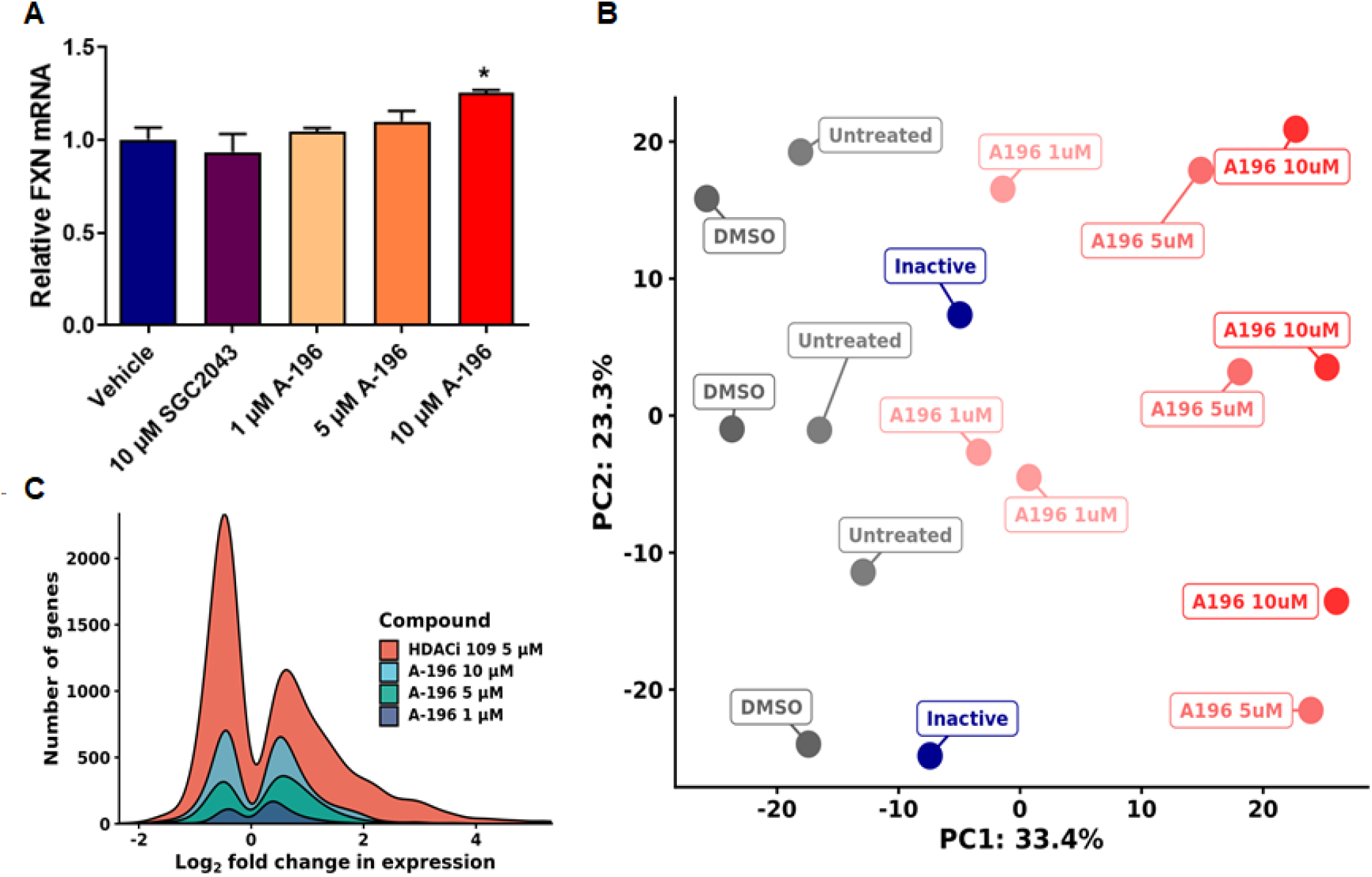
RNA Sequencing of primary FRDA fibroblast line GM04078 after A-196 treatment. **(A)** Relative abundance of *FXN* mRNA in A-196 and SGC2043-treated samples, demonstrating an increase in *FXN* expression. **(B)** PCA bi-plot of sequenced samples, illustrating a dose-dependent separation of untreated and A-196-treated samples along PC1. **(C)** Kernel density plot of significantly differentially expressed genes in all A-196-treated samples and HDACi 109-treated samples from Lai and colleagues (24). A-196 induces significantly lower transcriptional perturbation at all concentrations. Data are relative to the vehicle, a treatment of 6 days and are presented as mean ± SEM, n=3, one-way ANOVA followed by Bonferroni test *p<0.05.

We compared our transcriptomics results with previously published data reported from a histone deacetylase inhibitor (HDACi) (33) to measure genome-wide changes in expression. Genome-wide perturbations of expression between compound-treated and control samples were markedly lower in A-196-treated samples (A-196 1-10 μM: 193-1098 DEGs vs HDACi 5 μM: 3478 DEGs) (Figure 5C). In addition, the absolute magnitude of expression change by A-196 is lower compared to HDACi (A-196 1-10 μM: 45-67% vs HDACi 5 μM: 94%).

No pathways or gene sets were found to be significantly overrepresented within genes upregulated by A-196. In downregulated genes, collagen fibril organisation (GO:0030199) and the endoplasmic reticulum stress response (GO:0034976) are enriched, suggesting some perturbation of these pathways at high concentrations.

Targeting SUV-420 may promote a transition to an H4K20 monomethylated state, as observed by Schotta and colleagues (34). We examined the expression of seventeen key genes that represent affected pathways in this monomethylated state (Supplementary Table 2). Only two of these genes were differentially expressed using 10 μM A-196 (*CDK1* and *CCNB1*), with no significant perturbation at 5 μM and 1 μM. Combined with our genome-wide results, this suggests that A-196 is able to promote *FXN* expression with limited unwanted epigenetic effects.

### A-196 structural analogues increase *FXN* expression with improved potency

We sought to improve the potency of A-196 for increasing *FXN* expression through chemical structural modification (see Supplementary information). Information about the structure-activity relationship (SAR) of A-196 was already known, with certain key structural features identified as being critical for inhibitory activity (32). Examination of the crystal structure of A-196 bound to SUV4-20 H1 (Supplementary Figure 4) revealed potential avenues of exploration for synthetic modification. The cyclopentyl moiety occupies a hydrophobic pocket that could be further extended into, whilst the pyridyl group sits in a solvent-exposed region and potentially engages in a hydrogen-bonding interaction, which could be exploited. With this in mind, our efforts primarily focused on increasing lipophilic bulk at the cyclopentyl position, through either further extension into the pocket or increasing ring size. Alongside these modifications, a slight variation of the pyridyl group was also explored however not as exhaustively as with the upper-portion of A-196.

We treated the *FXN*-GAA-Luc cell line for six days with 5 μM of each new compound and found that four new molecules (compounds A3, A12, A14 and A15) increased FXN-Luc protein expression to levels comparable to that of A-196 (Figure 6A). These four compounds were also the most potent analogues synthesised, displaying the lowest IC_50_ values against SUV4-20 H1 (see Supplementary Table 3 and Supplementary Figure 5). Compounds A3 and A12 exert the strongest effect on FXN-Luc protein expression, and also show the greatest difference in potency between SUV4-20 H1 and SUV4-20 H2. Alongside the results of the siRNA mediated knockdown of SUV4-20 H1 and H2 (Figure 2A-D), this further suggests an enhanced role for SUV4-20 H1 in FXN expression, and may indicate an antagonistic role for SUV4-20 H2 inhibition.

**Figure 6.**
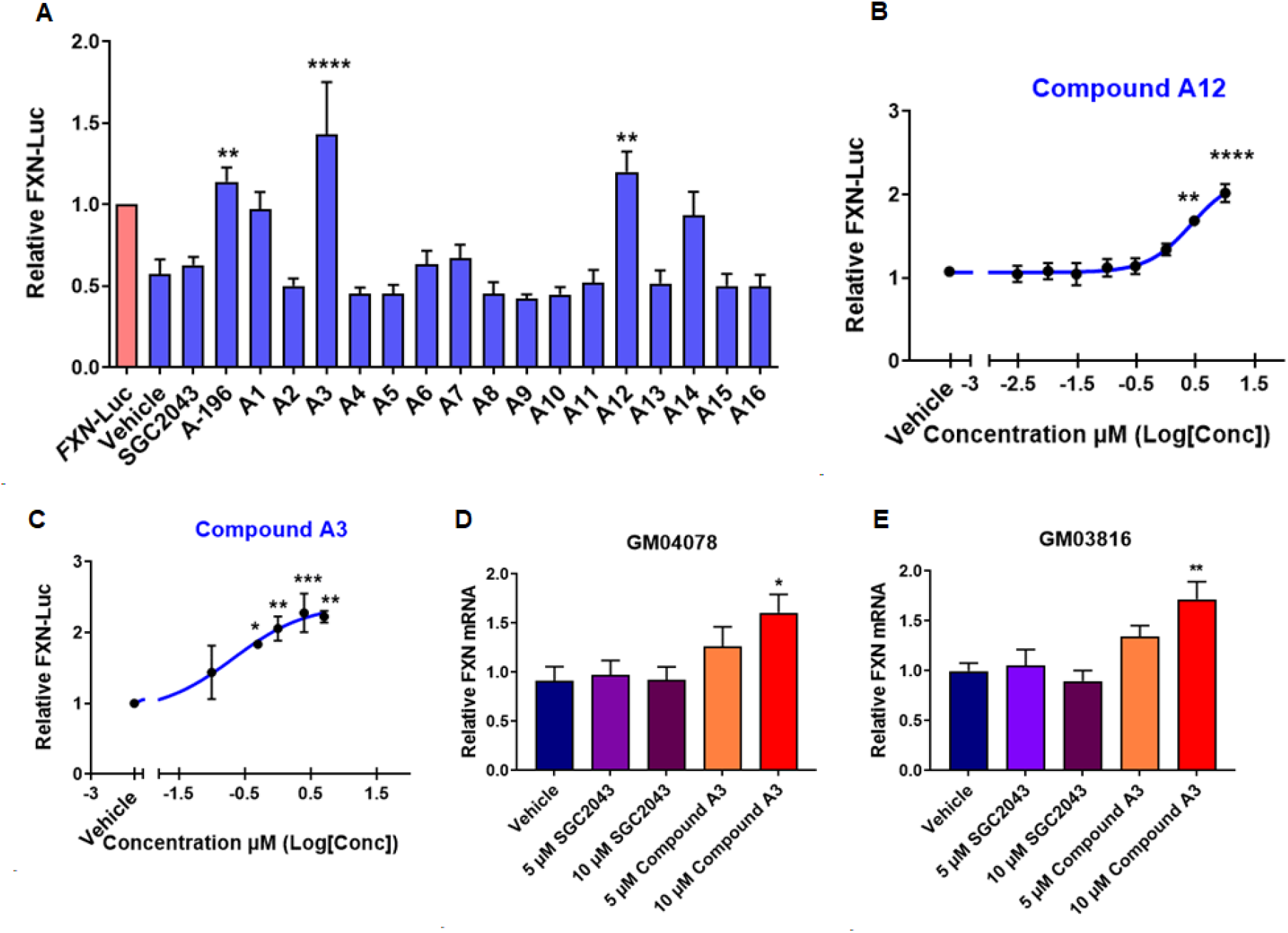
Medicinal chemistry synthesis of new A-196 derivatives capable of increasing frataxin protein expression. **(A)** Screen of A-196 derivatives using the *FXN*-GAA-Luc cell line, n=3 in triplicate **(B)** Concentration-response curve of compound A12 (EC_50_ 2.7 μM) in the line *FXN*-GAA-Luc, n=3 in triplicate **(C)** Concentration-response curve of compound A3 (EC_50_ 0.21 μM) in the line *FXN*-GAA-Luc, n=3 in triplicate **(D and E)** *FXN* mRNA expression of primary fibroblast GM04078 and GM03816 after treatment with compound A3, n=3 in duplicate. Data are relative to the vehicle, a treatment of 6 days and are presented as mean ± SEM, one-way ANOVA followed by Bonferroni test *p<0.05, **p<0.01, ***p<0.001, ****p<0.0001.

We next performed concentration-response assays using compounds A3 and A12 and confirmed a concentration-dependent increase in FXN-Luc expression after treatment with compounds A3 and A12 (Figure 6B and C). The estimated EC_50_ of compound A3 is 0.21 μM and of compound A12 is 2.7 μM, compared with 5.2 μM for the starting compound A-196. Finally, we treated patient-derived fibroblasts with compounds A3 and A12 for six days and found that compound A3 increased significantly the expression of frataxin mRNA (Figure 6D, E).

These SAR findings provide proof of concept that other active structural analogues of A-196 also increase *FXN* expression across cell lines and opens a new avenue for the potential discovery of clinical lead compounds.

## DISCUSSION

Our work builds on the intense research of recent years identifying the epigenetic mechanisms of *FXN* gene silencing in FRDA caused by the GAA repeat expansion in intron 1. Epigenetic silencing of the *FXN* gene is driven by several post-translational modifications of histones (13, 19, 24–31). Here we highlight the role of histone methylation in the regulation of *FXN* expression compared to the more widely-studied histone acetylation. We have identified and validated the histone methyltransferase SUV4-20 H1 as a novel therapeutic target for FRDA, first through pharmacological inhibition, and then siRNA-mediated gene knockdown, in a range of FRDA patient-derived primary cells.

Well-characterized libraries of probe compounds, such as the SGC epigenetic probe set screened here, allow researchers to link selective inhibition of a specific target with a biological response. Screening the SGC epigenetic modifiers allowed us to identify both new mechanisms of epigenetic gene regulation at the *FXN* locus and potential therapeutic targets for the treatment of FRDA. Interestingly, the screen of several inhibitors of chromatin regulatory proteins or domains (bromodomains, demethylases, and methyltransferases), highlighted the role of histone methylation in *FXN* repression. This suggests that by modulating the activity of some methyltransferases it is possible to increase the expression of the partially silenced frataxin gene in FRDA.

A-196 is a selective inhibitor of the SUV-420 methyltransferases (SUV4-20H1 and SUV4-20H2). It has been described to bind selectively to the SUV4-20 enzymes over 29 other methyltransferases, including protein arginine methyltransferases and DNA methyltransferases, a panel of chromatin binders and epigenetic readers, and 125 non-epigenetic targets, which include kinases, G-protein-coupled receptors (GPCRs), transporters and ion channels (32). Previous biochemical and co-crystallization analyses have shown that A-196 is a substrate-competitive inhibitor of both SUV4-20 H1 and H2 enzymes. In cells, this chemical probe induces a global decrease in di- and tri-methylation of H4K20, with a concomitant increase in mono-methylation (32). Our results show that this methyltransferase inhibitor produces a significant increase in *FXN* expression in FRDA patient-derived cells. In fibroblasts and lymphocytes, the up-regulation is by approximately 1.5 fold and in FRDA PBMCs by approximately two-fold. Asymptomatic carriers of the GAA expansion have only ~50% of the normal levels of frataxin (35) suggesting that even a modest increase in expression may be beneficial for FRDA patients.

The *FXN* gene carrying the disease-associated GAA repeat expansion exhibits increased H4K20me3 in the flanking regions of the GAA repeats (19) and decreased H4K20me1 downstream of the repeats (31). These two methylation marks are regulated by the chemical probe A-196 which has been shown to reduce global levels of H4K20me3 and H4K20me2 while increasing at the same time H4K20me1 (32). Although the siRNA-mediated knockdown of the targets we identify provides evidence of the importance of H4K20 methylation in the regulation of expression of *FXN,* histone lysine methylation and histone arginine methylation have not been extensively studied in FRDA, despite the fact that several reports describe histone methylation marks in the GAA expanded region (17).

One of the main concerns of epigenetic therapies is the potential broad biological effects that can result from the global inhibition of an epigenetic target. These effects may be caused by multiple genes that can be transcriptionally controlled by post-translational modifications in histone and non-histone proteins (36). A-196 demonstrated transcriptional activity that was dose-dependent and substantially reduced compared to a previously-reported HDACi (33). There was no evidence of specific pathway activation, suggesting that the transcriptional effects of A-196 do not result in significant changes in cellular function. This activity is most likely due to SUV4-20 inhibition: at 1 μM and 10 μM, A-196 has been shown to selectively inhibit SUV4-20 (27). No other protein lysine methyltransferases, including the other methyltransferase modifying H4K20, namely SET8, or those that use H3K4, H3K9, H3K27, and H3K79 as substrates, were found to be inhibited by A-196 at that concentration. This high level of specificity is of great importance and selective inhibitors of specific methyltransferases, if found to modify certain pathologies, may have therapeutic potential with possibly fewer side effects than other epigenetic targets such as HDACs.

Previous studies of FRDA have focussed on achieving chromatin remodelling by HDACi (24, 26, 43, 44, 30, 33, 37–42). As opposed to acetyltransferases, lysine methyltransferases have high specificity, modifying usually one single lysine on a single histone. These modifications can either lead to activation or repression of transcription (45). Furthermore, at least 50 non-histone proteins have been reported to be HDAC substrates, including several transcription factors (RUNX3, p53, c-Myc, nuclear factor kappa-light-chain-enhancer of activated B cells), chaperones (HSP90), signalling mediators (Stat3 and Smad7) and DNA repair proteins (Ku70) (36) which may contribute to potentially-detrimental genome-wide effects of HDAC inhibition in patients.

In contrast, lysine methyltransferases have been shown to modify only a few non-histone substrates (46, 47). Amongst them, p53 is the most common modified protein, since it can be methylated by the methyltransferases Set7/9 (K372), SMYD2 (K370), G9a (K373), GLP (K373) and SET8 (K382) (46, 48). To the best of our knowledge, amongst the methyltransferases responsible for the methylation of H4K20, only SET8 (H4K20me1) is able to methylate non-histone proteins such as p53 and proliferating cell nuclear antigen (PCNA) (47). To date, no studies have reported that either of the SUV4-20 methyltransferases is able to methylate non-histone proteins, suggesting their selectivity for histones. Moreover, it has been shown that this family of enzymes preferentially methylate histone H4K20 on nucleosomes rather than free histones (20), narrowing down in this way, even more, the epigenetic modification of SUV4-20 genome-wide. The aforementioned characteristics of lysine methyltransferases may be the reason why A-196 used at all tested concentrations, modifies the expression of fewer genes compared to HDACi (33).

Although A-196 is highly selective, a caveat for the inhibition of SUV4-20 may be the possible genome-wide transition to an H4K20 mono-methylated state. Schotta and colleagues reported in mice that a chromatin-wide transition to H4K20me1 impairs genome integrity and programmed DNA rearrangements (34) and that the complete loss of both SUV4-20 enzymes in *SUV4-20h1*^*−/−*^ *SUV4-20h2*^*−/−*^ double-null mice resulted in ablation of nearly all H4K20me3 and H4K20me2, a change incompatible with embryonic development. A genome-wide transition to a mono-methylated H4K20 state led to increased sensitivity to damaging stress, with a mechanism that is dependent on inefficient DNA double-strand break repair and consequent chromosomal aberrations. B cells lacking SUV4-20 were shown to be defective in immunoglobulin class-switch recombination, a process required for antibody isotype diversification. Chemical inhibition of SUV4-20 may, however, be modulated to bypass complete inhibition of these methyltransferases, thus avoiding any aberrant transition of H4K20 methylated states. In support of this, we did not see evidence from our transcriptomic data of A-196 inducing these states and find that unwanted epigenetic activity is even further reduced at the lower doses of 5 μM and 1 μM.

Improving the potency of A-196 will likely be a strategy to avoid extensive modification of the epigenome, as it will permit the use of lower doses and limit the risks of off-target effects. Chemical structural modification of A-196 produced compound A3, which according to our estimated EC_50_, is 24 times more potent than A-196 at inducing *FXN* expression (Figure 1C and Figure 6C). New compounds that selectively target SUV4-20 H1 may be important for potentially treating FRDA patients and as important tools to further explore the role of this methyltransferase in other disease states. Our data show that structural modification of A-196 can generate new compounds with the ability to increase *FXN* expression. This increase was observed when using the *FXN*-GAA-Luc cell line as well as primary fibroblasts, an indication that *FXN* increase in expression is not cell-dependent or limited to a particular cell line.

## MATERIALS AND METHODS

### Materials

#### Cell lines

Compound screening was performed using the *FXN*-GAA-Luc reporter cell line with the *FXN*-Luc cell line used as a reference for normal frataxin levels. Both cell lines were built in a HEK293 background as previously described (49) and carry the entire 80 kb human *FXN* locus (exons 1–5b of *FXN*) with an insertion of the luciferase gene in exon 5a and contain either 6 GAA repeats (*FXN*-Luc) or ~310 GAA repeats (*FXN*-GAA-Luc) in intron 1 of the *FXN* gene. The patient-derived cells used were patient-derived fibroblast lines GM04078 (from an FRDA clinically affected individual carrying alleles with 541 and 420 GAA repeats) and GM03816 (from an FRDA clinically affected individual carrying alleles with 330 and 380 GAA repeats). Patient-derived lymphoblastoid cell lines GM158850 (from an FRDA clinically affected individual carrying alleles with 650 and 1030 GAA repeats) and GM016220 (from an FRDA clinically affected individual carrying two alleles each with 460 GAA repeats) were also utilized.

### Methods

#### Cell culture

The HEK293 reporter cell lines were cultured in Dulbecco’s modified Eagle’s medium (DMEM) – high glucose supplemented with 10% fetal bovine serum (FBS), 2 mM L-glutamine, 100 U/ml penicillin/streptomycin and 100 μg/ml Hygromycin B (Life Technologies) (complete DMEM medium). Fibroblasts lines were obtained from Coriell Institute (USA) and cultured in Minimum Essential Medium (MEM) supplemented with 15% FBS, 1% MEM Non-essential Amino Acid Solution (100X Gibco) and 100 U/ml penicillin/streptomycin. Lymphoblastoid cell lines were also obtained from Coriell Institute (USA) and cultured in RPMI medium with 15% FBS, 2 mM L-glutamine and 100 U/ml penicillin/streptomycin. Peripheral blood mononuclear cells (PBMCs) were isolated according to Smith et al (50) and cultured in RPMI medium supplemented with 10% FBS, 2 mM L-glutamine and 100 U/ml penicillin/streptomycin.

### Luciferase assay

For assessment of frataxin-luciferase protein expression, cells were washed with PBS and lysed directly on the plate on ice using the Cell Culture Lysis Reagent from the Luciferase Assay System (Promega#E1500). Lysates were transferred to microcentrifuge tubes, vortexed for 15 sec and centrifuged at 12.000 x g for 15 sec. The supernatant was decanted into a microcentrifuge tube and 25 μl was loaded in a white opaque 96-well microplate (PerkinElmer). Luciferase expression was determined by measuring luminescence with a PHERAstar FSX microplate reader (BMG LABTECH) equipped with injectors pumps, where the injection of 100 μl of Luciferase Assay Reagent was loaded to each sample before measurement. Data were normalized to total protein concentration as determined by BCA protein assay (ThermoFisher).

### Screening the Structural Genomics Consortium epigenetic chemical probes library

*FXN*-GAA-Luc cells were seeded in 96-well plates at a density of 5 x 10^3^ in duplicate and allowed to recover overnight before treatment with the SGC epigenetic probes set. The library was obtained directly from the Structural Genomics Consortium. Concentrations and incubation times are as described in Supplementary Table 1. The Luciferase assay for frataxin-luciferase protein expression was performed as described above.

### Western blot assay

Cells were washed with PBS and lysed directly on the plate on ice with RIPA buffer [(50 mM, Tris, pH 8, 150 mM NaCl, 2 mM EGTA, 0.5% sodium deoxycholate, 1% igepal 630, 0.1% sodium dodecyl sulphate (SDS) with protease inhibitors (Complete Mini, EDTA-free, Roche)]. The lysates were then transferred to a collection tube and sonicated for a few seconds prior centrifugation for 15 min at 300 x g and 4°C. Protein concentration was determined using a BCA protein assay (ThermoFisher). Samples were then reduced in Laemmli buffer and incubated for 5 min at 95°C. Thirty micrograms of protein were resolved for 50 min at 200V on 8%-15% SDS-polyacrylamide gel electrophoresis. Following wet transfer on a PVDF membrane (BioRad), samples were incubated with the following antibodies: mouse monoclonal anti-frataxin (Abcam ab113691, 1/1000), rabbit polyclonal anti-G9a (Cell signaling #3306 1/5000), rabbit monoclonal anti-EZH1 (Cell signaling #42088 1/1000), rabbit monoclonal anti-EZH2 (Cell signaling #5246 1/2500), rabbit polyclonal anti-DOT1L (Abcam ab72454, 1/1000), rabbit polyclonal anti-H4K20me1 (Abcam ab9051, 1/500), rabbit polyclonal anti-H4K20me2 (Abcam ab9052, 1/2000), rabbit polyclonal anti-H4K20me3 (Abcam ab9053, 1/2000), rabbit polyclonal anti-H4 (Abcam ab10158, 1/2000), and HRP conjugated anti-Beta-Actin (Abcam ab49900 1/25000). Protein quantities were analysed using Image Lab software.

### siRNA-mediated down-regulation of targets

*FXN*-GAA-Luc cells were counted with an automated cell analyser NucleoCounter^®^ NC-250TM (ChemoMetec) and seeded in a 12 well poly-L-lysine coated plate (12000 cells/well) in complete DMEM medium (see above) minus antibiotics. Cells were then allowed to recover overnight. Next day cells were rinsed in Opti-MEM (GibcoTM 31985047) prior to the liposomes delivery. Equal volumes of siRNA and RNAiMAX transfection reagent (GibcoTM 13778075) were mixed and allowed to form complexes for 20 minutes. The mix was then added to the cells drop-by-drop and incubated for six hours. Subsequently, the medium was supplemented with FBS and next day, the medium was changed to complete DMEM medium (see above) minus antibiotics. Six days post-transfection cells were collected for western blot, luciferase and qRT-PCR assays. siRNAs were purchased from Dharmacon as SMARTpool/ON-TARGETplus as follow: SUV4-20 H1 (#51111), SUV4-20 H2 (#84787), EZH1 (#2145), EZH2 (#2146), DOT1L (#84444), and non-targeting (#D-001810-10).

### Adenylate kinase assay

The adenylate kinase assay (ToxiLightTM bioassay kit LT07-217 Lonza) to assess toxicity was performed following the instructions of the manufacturer. Briefly, 5 μl of the media where cells were cultured was transferred to a 384 Greiner LUMITRACTM white plate and allowed to reach room temperature (RT). Twenty-five microliters of the adenylate kinase detection reagent were then added to each well and incubated for five minutes. Finally, the plate was read in the PHERAstar FSX microplate reader (BMG LABTECH).

### qRT-PCR

Total mRNA was extracted using the RNeasy Mini Kit (Qiagen) and then treated with RNase-free DNase (Qiagen). One microgram of RNA was used to synthesize cDNA using random primers (Life Technologies) and SuperScript III Reverse Transcriptase (Life Technologies) in a 20-μl reaction volume. qPCR was carried out using primers which can be found in the supplementary information. Data were normalized to HRPT1 and analysed by the 2^−ΔΔ^_Ct_ method (51).

### AlphaLISA assay to quantify human frataxin

The AlphaLISA human frataxin detection kit (PerkinElmer, catalogue # AL322HV/C/F) was scaled down to a white opaque 384-well plate format (PerkinElmer) and adapted from the manufacturer’s recommendations. Briefly, an hFXN analyte standard dilution was prepared alongside the samples using the AlphaLISA immunoassay buffer, to which the biotinylated anti-hFXN Antibody was added for one hour at RT. AlphaLISA anti-hFXN acceptor beads were added for an additional hour at RT, after which streptavidin donor beads solution were put on for a further 2 hours at RT. Finally, the plate was read using a PHERAstar FSX microplate reader (BMG LABTECH) at a wavelength of 615 nm. Data were normalized for protein quantity using the BCA assay.

### RNA-Sequencing

Total RNA from primary fibroblasts was extracted using the RNeasy Mini Kit (Qiagen), including DNAse I treatment to remove genomic DNA contamination. RNA integrity was determined using the Agilent RNA 6000 Pico kit on a Bioanalyzer. One hundred ng of each sample (measured using the Quant-iT RiboGreen RNA Assay Kit) was submitted for library preparation and sequencing at the Oxford Genomics Centre. Twenty-four strand-specific libraries were prepared from GM04078 and the healthy control line GM03440 using the NEBNext Ultra II mRNA kit following manufacturer’s instructions. Sequencing was performed on an Illumina HiSeq 4000 across two lanes, generating 25-33 million read pairs per sample. Reads were pseudoaligned to Ensembl GRCh38 (version 98) cDNA reference using Kallisto (version 0.46.0), with 86-89% alignment. Transcript abundance estimates were summarised to gene-level counts using Tximport (version 1.12.3). Differential gene expression analysis was performed using DESeq2 (version 1.24.0), excluding genes with fewer than 10 counts average across all samples. An FDR cut-off of ≤0.01 was used to call differential expression. Principal component analysis was performed using prcomp in R (version 3.6.1). Two GM03440 samples and one GM04078 sample were classed as outliers due to isolated clustering from technical replicates on a PCA bi-plot and were therefore excluded from further analysis (Supplementary Figure 6).

Pathway enrichment analysis was performed using gprofiler2 (version 0.1.8), querying GO Biological Process, KEGG and REACTOME databases. Gene sets larger than 350 were excluded for interpretability. All genes considered for differential expression were used as background and an FDR cut-off of 0.01 was used.

For comparison between our transcriptomic data and the work by Lai and colleagues (33), raw fastq files were obtained from Sequence Read Archive (SRA) accession PRJNA495860 and processed as described above. The number of replicates per condition was kept constant to minimise differences in statistical power.

## DATA AVAILABILITY

RNA sequencing data from this publication are available at Gene Expression Omnibus (GEO) (https://www.ncbi.nlm.nih.gov/geo/query/acc.cgi?acc=GSE145115) under accession number GSE145115.

## ACKNOWLEDGEMENT

We would like to thank Dr Ana M. Silva and Dr Michele M.P. Lufino for the development of the *FXN*-GAA-Luc and the *FXN*-Luc reporter cell lines, and for advice at the start of this study. We also thank the Oxford Genomics Centre at the Wellcome Centre for Human Genetics (funded by Wellcome Trust grant reference 203141/Z/16/Z) for the generation and initial processing of the sequencing data.

